# The role of two *GLYCOALKALOID METABOLISM* genes in α-tomatine biosynthesis and basal defense in tomato

**DOI:** 10.64898/2026.02.16.705890

**Authors:** Yaohua You, Aishwarya Balaji, Andrea Lorena Herrera Valderrama, Marie-Emma Denarie, HM Suraj, Miguel Ramirez Gaona, Katharina Hanika, Francel Verstappen, Iris F. Kappers, Jan A.L. van Kan

## Abstract

Steroidal glycoalkaloids and saponins are plant cholesterol-based steroid metabolites with antimicrobial activities and potential pharmacological value. The saponin uttroside B from black nightshade (*Solanum nigrum*) plays an important role in defense against herbivorous insect and exhibits anti-hepatocellular carcinoma activity. The tomato (*S. lycopersicum*) glycoalkaloid α-tomatine has been studied because of its antinutritional effects, however, its role in protecting plants from fungal pathogens remains understudied. The biosynthetic pathway of α-tomatine involves multiple clustered genes designated as *glycoalkaloid metabolism* (*GAME*) genes. In this study, we generated single knockout mutants of *SlGAME4* and *SlGAME2* by CRISPR/Cas9-based genome editing. The *SlGAME4* mutants did not accumulate glycoalkaloids but instead redirected resources towards steroidal saponin (uttroside B) synthesis. *SlGAME2* mutants contained unaltered α-tomatine contents indicating that the *SlGAME2* gene, previously reported to catalyze the transfer of xylose to β_1_-tomatine, is not involved in α-tomatine biosynthesis. Infection assays with four fungal tomato pathogens demonstrated that the *SlGAME4* mutant plants were slightly more susceptible to *Botrytis cinerea*, but equally susceptible to the other three fungi. Up-regulation of α-tomatine-responsive genes in *B. cinerea* was observed during infection on *SlGAME4* mutant tomato, as well as on *S. nigrum* suggesting that uttroside B induces a fungal transcriptional response similar to α-tomatine. Furthermore, we observed that tolerance mechanisms to plant saponins mediated by glycosyl hydrolases and a glycosyltransferase contribute to virulence of *B. cinerea* on *SlGAME4* mutant plants and *S. nigrum*. This indicates that also uttroside B contributes to defense against fungal pathogens and can be detoxified by *B. cinerea*.

## Introduction

Tomato (*Solanum lycopersicum*) from the Solanaceae (nightshade) family is an economically important vegetable crop and serves as a rich source of nutrition worldwide. The yield of tomato can be affected by pathogenic fungi such as grey mold (*Botrytis cinerea*), leaf mould (*Cladosporium fulvum*), Verticillium wilt (*Verticillium dahliae*), early blight (*Alternaria solani*) or by oomycetes such as late blight (*Phytophthora infestans*), as well as by herbivorous insects (Arie et al., 2007; Blancard, 2012; Panthee and Chen, 2010; Nowicki et al., 2012). It highlights the necessity to study basal resistance traits of tomato, in particular its reservoir of endogenous antimicrobial metabolites with defensive roles (Bednarek et al., 2012).

Steroidal glycoalkaloids (SGA) are a subgroup of saponins constitutively produced by plant species in the Solanaceae and Liliaceae families (Cárdenas et al., 2015). They not only possess antinutritional properties (bitterness) but are also considered phytoanticipins protecting the plants from attack by pathogens and herbivores due to their high concentration and broad-spectrum antimicrobial as well as insecticidal activities (Sandrock and VanEtten, 1998; Sun et al., 2021; Zhao et al., 2021). α-tomatine is the major SGA in tomato and accumulates in vegetative tissues and green fruit to concentrations exceeding 1 mM (Kozukue et al., 2004; You and van Kan, 2021). The biosynthetic pathway of SGAs starts from the precursor cholesterol and is mediated by enzymes encoded by *GLYCOALKALOID METABOLISM* (*GAME*) genes (Itkin et al., 2013). The GAME4 gene product, a cytochrome P450 protein, catalyzes the first dedicated step from furostanol towards SGAs (Itkin et al., 2013). The intermediate alkaloid tomatidine, the aglycone of α-tomatine, does not have antimicrobial activity but is toxic to plants (Ökmen et al., 2013). This phytotoxicity can be mitigated by four consecutive glycosylation steps (catalyzed by GAME1, GAME17, GAME18 and GAME2), ultimately resulting in production of α-tomatine (Itkin et al., 2013). During fruit ripening, α-tomatine is exported from the vacuole to the cytosol by the tonoplast transporter GORKY and converted into a less bitter, non-toxic SGA, named esculeoside A (Cárdenas et al., 2015; Kazachkova et al., 2021).

The toxicity of α-tomatine to fungi is attributed to the disruption of fungal plasma membranes through complexing with 3β-hydroxysterol (Steel and Drysdale, 1988; You and van Kan, 2021). Enzymatic detoxification of α-tomatine has been reported in many tomato pathogens, and the most studied mechanism involves secreted glycosyl hydrolases (GH) referred to as ‘tomatinase’. Tomatinase activity in bacteria and fungi that are pathogenic on tomato has been reported for glycosyl hydrolases from three distinct families (GH10, GH3 and GH43) (You and van Kan, 2021; You et al.,2024). Besides hydrolytic detoxification, *B. cinerea* also possesses multiple non-degradative mechanisms for tolerance to α-tomatine which are mediated by proteins involved in fungal membrane repair and modification (You et al.,2024).

The constitutive presence of antimicrobial metabolites makes an important contribution to basal plant defense (Osbourn et al., 1996; Zaynab et al., 2021). The most compelling evidence comes from saponin-deficient mutants of *Avena strigosa* (wild diploid oat), that exhibited compromised resistance to fungi that normally cannot infect oat, such as *Gaeumannomyces graminis* var. *tritici, Fusarium culmorum* and *F. avenaceum* (Papadopoulou et al., 1999). In a study with 23 fungi, Sandrock and VanEtten (1998) observed that seven taxa that are non-pathogenic on tomato were all sensitive to α-tomatine, while 14 out of 16 tomato pathogens were tolerant to it. We recently reported that *B. cinerea* isolate M3a from grape is sensitive to α-tomatine and could barely colonize tomato leaves, and that its virulence was enhanced by overexpression of genes that confer tolerance to α-tomatine (You et al., 2024). Here we describe the generation of knockout mutants in the tomato *SlGAME4* and *SlGAME2* genes via CRISPR/Cas9 and report the effects of these deletions on saponin profiles and on interactions with fungal pathogens.

## Results

### CRISPR/Cas9-mediated mutagenesis of *SlGAME4* and *SlGAME2* genes in tomato

We selected the *SlGAME4* and *SlGAME2* genes for mutagenesis for the following reasons. The *GAME4* gene product catalyzes the first dedicated step in the alkaloid biosynthetic pathway, using furastonol as substrate (Itkin et al., 2013; Grzech et al., 2025). Inactivating this gene would abolish synthesis of all SGAs. By contrast, the *GAME2* gene encodes a glycosyl transferase that was reported to catalyze the final step in α-tomatine synthesis, the transfer of xylose to β_1_-tomatine (Itkin et al.,2013). Inactivating this gene would result in the accumulation of β_1_-tomatine, which is non-toxic to fungi (Quidde et al., 1998).

CRISPR/Cas9 genome editing was employed to generate single knockout (KO) mutants of *SlGAME2* and *SlGAME4* with four sgRNAs targeting the open reading frame of each gene, in the background of *S. lycopersicum* cv. Moneymaker (MM). A high incidence of biallelic mutations was observed in the T0 generation of primary transformants: six out of seven *SlGAME2*-KO lines and all six *SlGAME4*-KO lines tested carried mutations in both alleles of the target genes. Deletions ranging from 1 bp to 1370 bp were detected in both *SlGAME2* and *SlGAME4* in the T0 generation and were inherited by T1 plants. **Figure 1** illustrates positions and sizes of deletions in each gene from four independent homozygous T1 mutants that were analyzed in detail. In most cases, deletions were near to the target sequences of either of the sgRNAs. However, the *SlGAME4*-KO line #6-7 contained a 617 bp deletion that starts 226 bp downstream of the predicted cleavage site of sgRNA1 and ends 229 bp downstream of the predicted cleavage site of sgRNA2.

**Figure 1.**
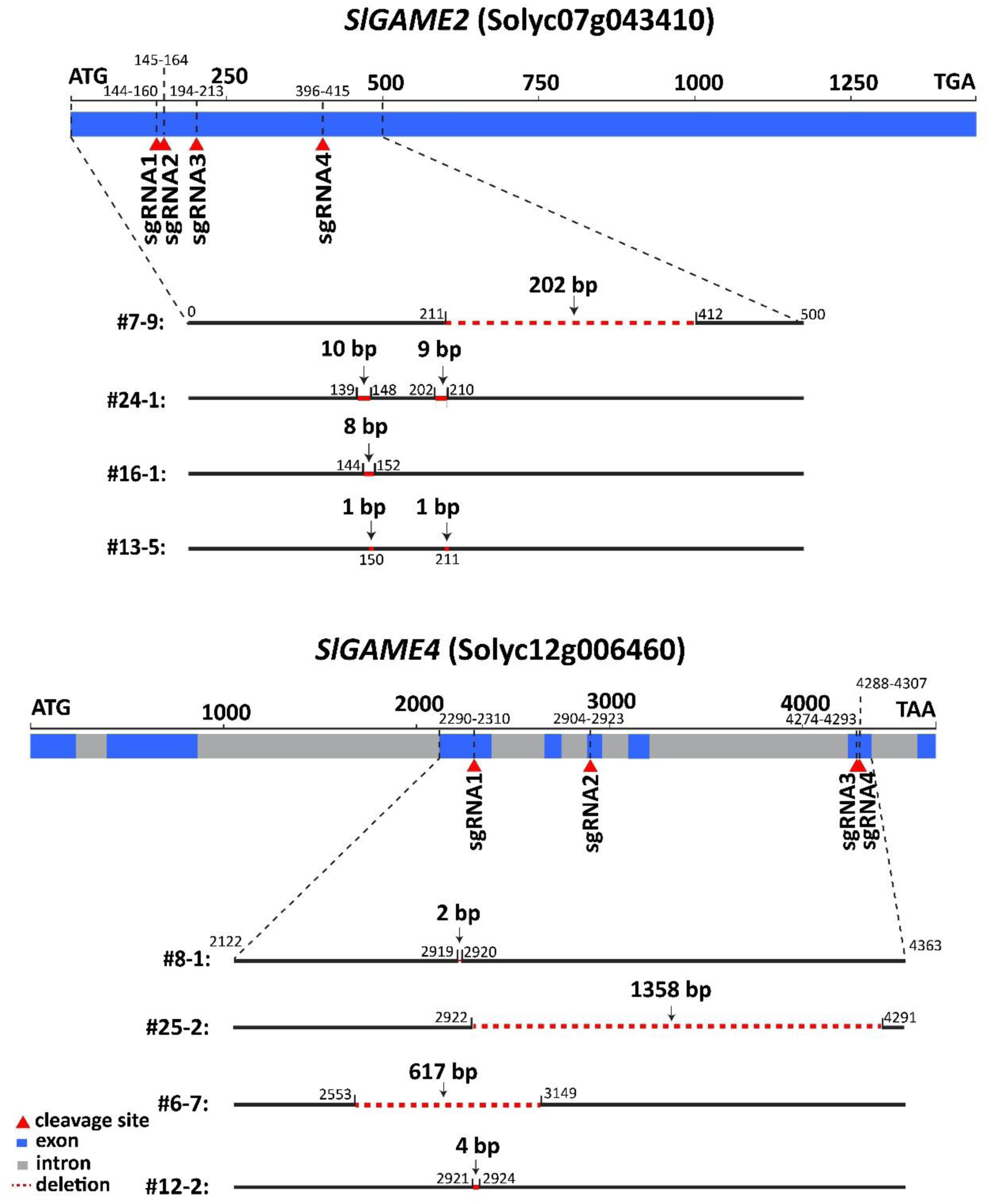
Scheme of CRISPR/Cas9-mediated KO of *SlGAME2* and *SlGAME4*. Positions of sgRNAs and their predicted cleavage sites are indicated by coordinates and red arrows, respectively. Deletions are illustrated by red dotted lines, the coordinates of the deleted nucleotides are provided, starting counting from the start codon.

### Analysis of tomatidine and α-tomatine contents in tissues of KO plants

Relative concentrations of α-tomatine and tomatidine were analyzed in young and mature leaves, stems and roots sampled from *SlGAME2*-KO, *SlGAME4*-KO and wild-type MM plants. α-Tomatine and tomatidine accumulation were abolished in *SlGAME4*-KO plants with only trace amounts of α-tomatine and tomatidine detected (**Figure 2**), possibly by cross-contamination from previous runs in LC-QqQ-MS. Strikingly, all four independent *SlGAME2*-KO lines produced α-tomatine at concentration similar to the MM recipient (**Figure 2**). Among the tissues sampled, young leaves contained by far the highest concentration of α-tomatine but not of tomatidine. The roots contained around 10 times less α-tomatine than young leaves but accumulated the highest content of tomatidine as compared to other tissues, with almost equal levels of α-tomatine and tomatidine (**Figure 2**). Stem tissues contained the lowest concentrations of both α-tomatine and tomatidine.

**Figure 2.**
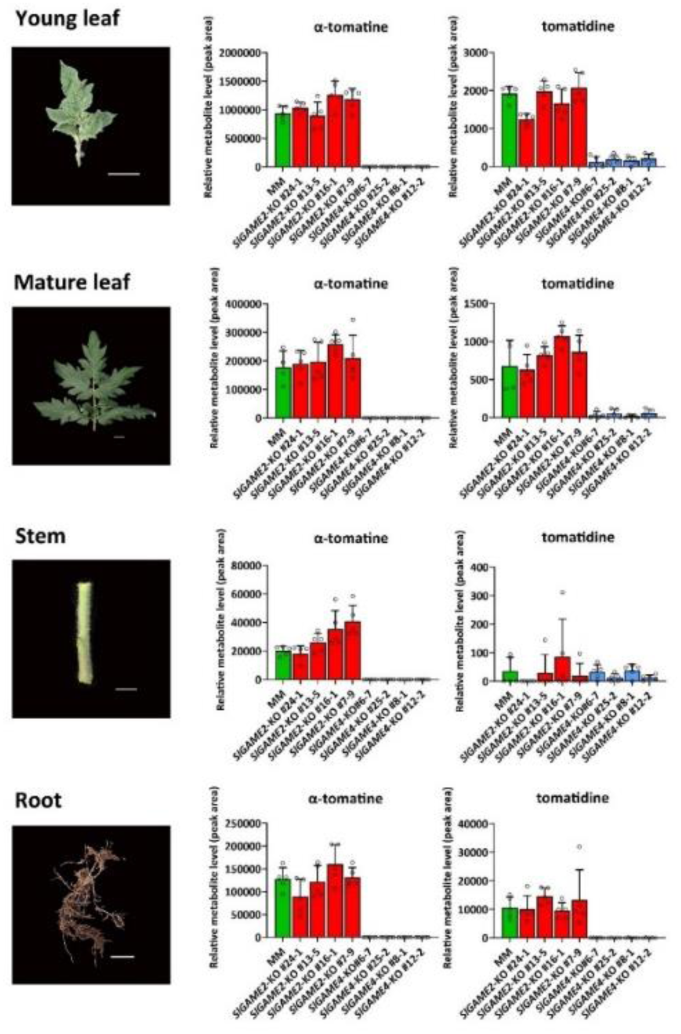
Content of α-tomatine and tomatidine in tissues of *SlGAME2*-KO, *SlGAME4*-KO and MM plants. The numbers underneath the columns indicate plant genotypes tested. Error bars are standard error of mean (SEM) of five biological replates. Scale bars in the images indicate 2 cm.

We analyzed the saponin profiles in *SlGAME4*-KO plants in more detail using LC-MS2. Leaves from four separate *SlGAME4*-KO mutant genotypes contained no detectable α-tomatine but instead accumulated uttroside B, while wild type MM leaves contained α-tomatine but no uttroside B **(Supplementary Figure 1)**. Fruit from MM and two independent *SlGAME4*-KO mutant genotypes was analysed, using green and red ripe fruit from three individual plants per genotype (**Figure 3)**. MM green fruit contained a high level of α-tomatine (compound 3), which in red ripe fruit was converted into esculeoside A (compound 1), as well established (reference). *SlGAME4*-KO mutant fruit neither contained α-tomatine nor esculeoside A but instead accumulated uttroside B (compound 4) and a compound annotated as uttroside B + pentose (compound 2). The ratio between compounds 4 and 2 decreased upon ripening in the GAME4-KO #12-2 line, while it remained similar in the GAME4-KO #25-2 line.

**Figure 3.**
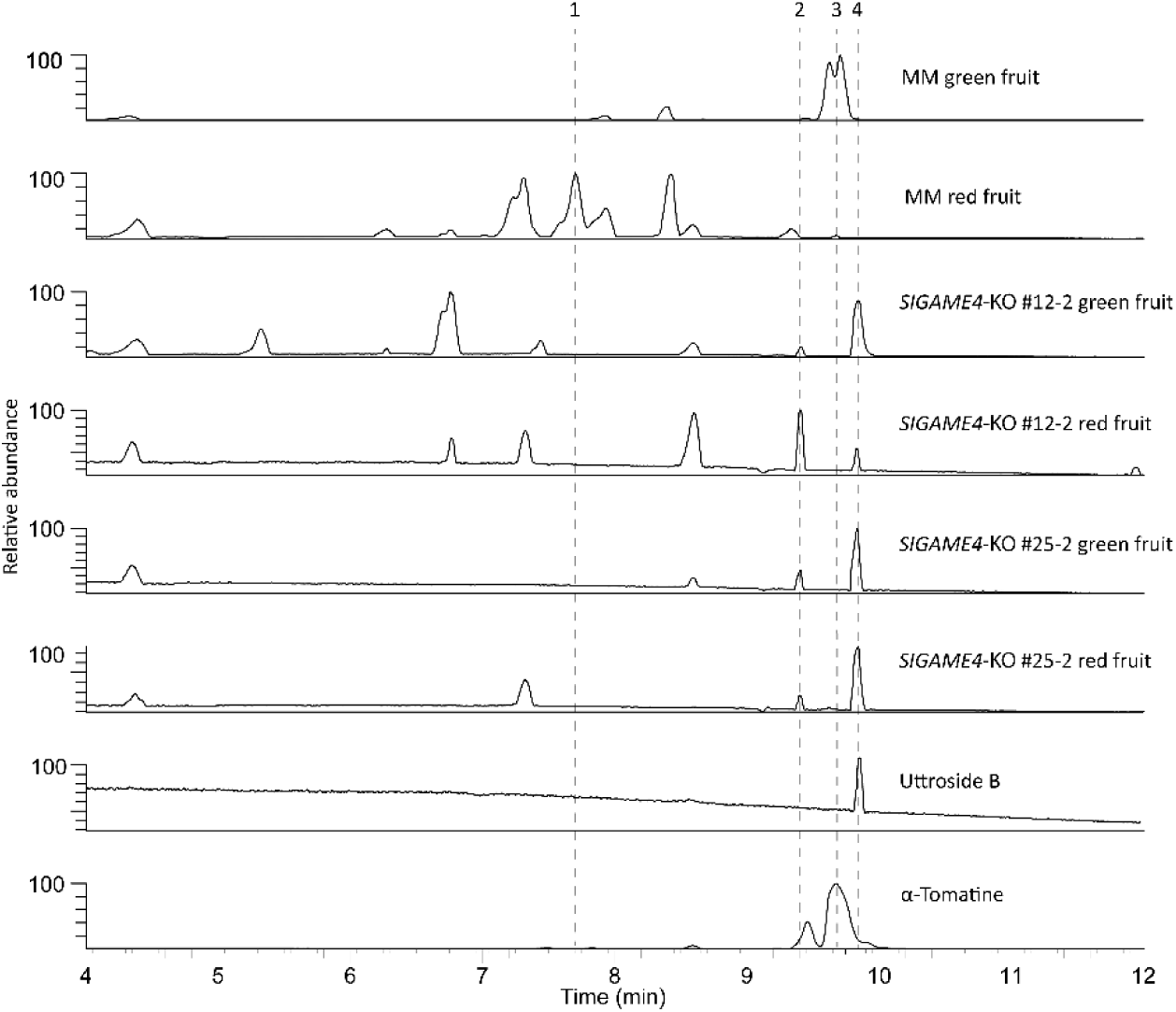
Metabolite analysis in green and red ripe fruit from wild type MM and *SlGAME*4 mutant tomato. Samples are labeled on the right hand side of the graph. Dotted lines mark peaks of the following metabolites: 1. esculeoside A (m/z 1270.6, RT= 7.7 min); 2. uttroside B + pentose (m/z 1329.6, RT = 9.4 mins); 3. α-tomatine (m/z 1034.55, RT= 9.7 min); 4. uttroside B (m/z 1197.59, RT= 9.8 min).

### Analysis of expression levels of α-tomatine biosynthetic genes in tomato

We analyzed the expression levels of several *GAME* genes, reported by Itkin et al. (2013), in young leaves, undeveloped leaves, mature leaves and stems in MM as well as in the wild tomato *Solanum habrochaites* LYC4 (**Supplementary Figure S2**). Remarkably, expression of *GAME2* was barely detected in MM by qRT-PCR. In LYC4, *GAME2* transcription was mainly detected in young and undeveloped leaves.

### Expression profile of fungal α-tomatine-responsive genes during plant infection

Transcriptional up-regulation of multiple α-tomatine-responsive genes was first demonstrated during *in vitro* growth of *B. cinerea* in the presence of α-tomatine (You et al., 2024). The genes *BcTom1* and *BcGT28a* play important roles in tomato infection and their expression was induced upon inoculation on tomato leaves, but not on *Nicotiana benthamiana* as this species does not accumulate α-tomatine (You et al., 2024). We analyzed the transcript levels of nine α-tomatine-responsive genes during infection on leaves of *SlGAME4*-KO plants by RT-qPCR. Despite the absence of α-tomatine in this plant, up-regulation of the genes was observed from 24 hpi onwards (**Figure 4**), which coincided with the onset of plant cell death induction and development of necrotic lesions.

**Figure 4.**
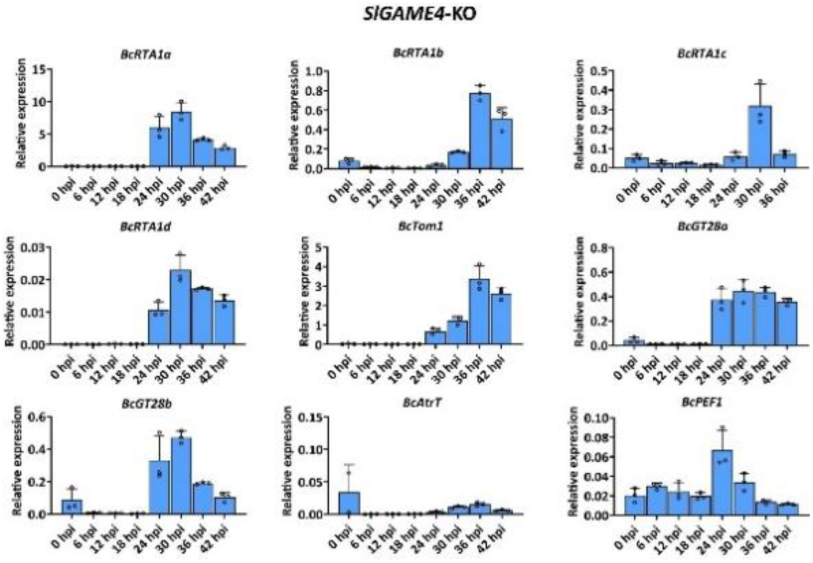
Relative expression of α-tomatine-responsive genes in *B. cinerea* during infection on *SlGAME4*-KO #6-7 leaves. Error bars are standard error of mean (SEM) of three biological replates.

### Susceptibility of *SlGAME4*-KO plants to different tomato pathogens

To assess the influence of the modification of saponin composition on disease susceptibility, we inoculated *SlGAME4*-KO plants with different tomato pathogens including the necrotrophic fungi *B. cinerea* and *Alternaria solani*, the soil-borne vascular pathogen *Verticillium dahliae* and the biotrophic leaf mould *Fulvia fulva*. Three independent *SlGAME4*-KO mutant plants showed a slight increase in susceptibility to *B. cinerea* but were not significantly altered in susceptibility to the other three fungi (Figure 5).

**Figure 5.**
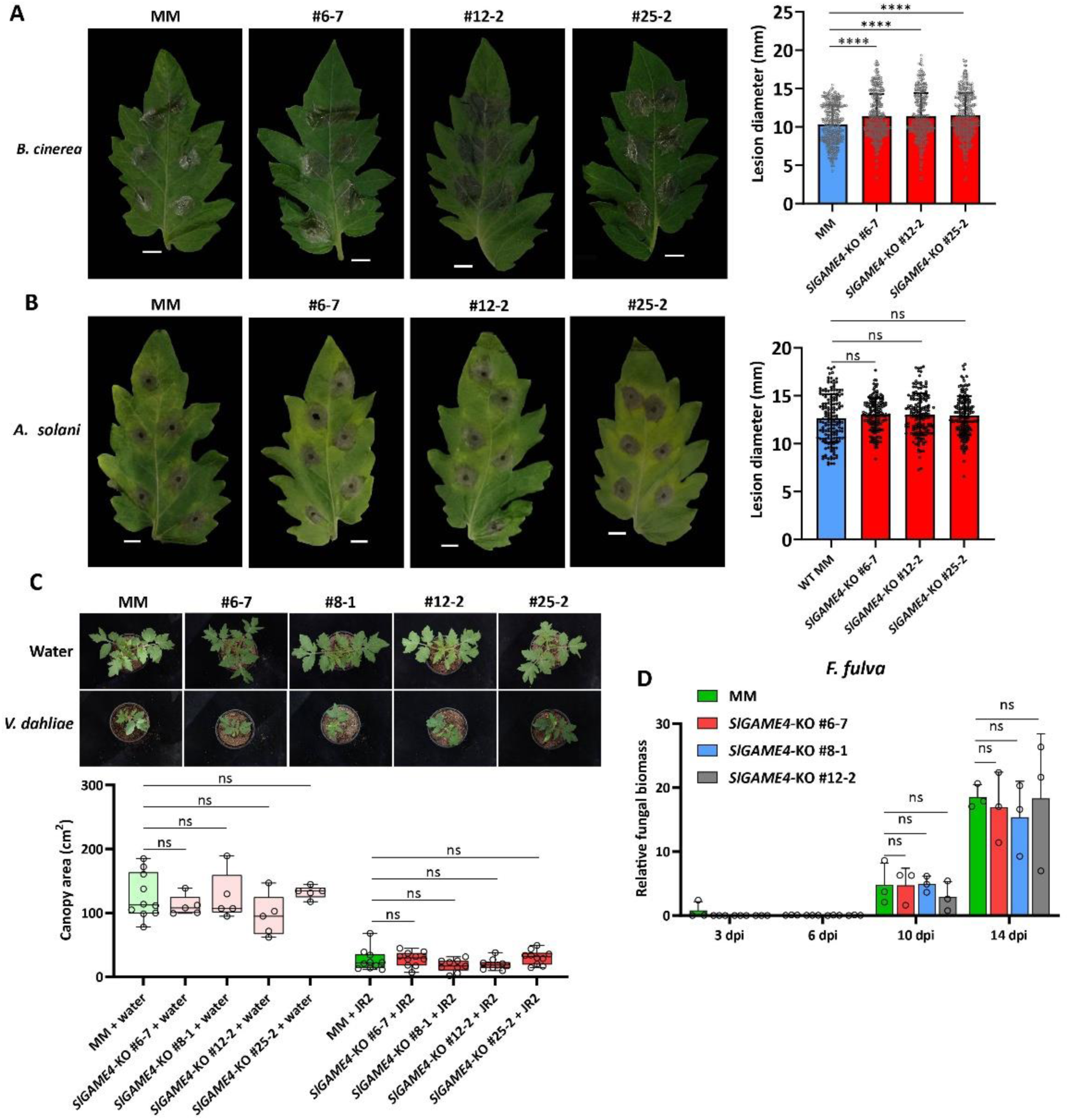
Susceptibility to fungal pathogens of *SlGAME4*-KO mutant plants. Lesion diameters of *Botrytis cinerea* (A) and *Alternaria solani* (B) after inoculation on leaves of *SlGAME4*-KO were measured at 3 dpi and 5 dpi, respectively. (C), canopy areas (cm^2^) of tomato seedlings inoculated at the roots with water or with *Verticillium dahliae* strain JR2 at 15 dpi. Canopy areas were calculated with ImageJ, from photographs that were taken from above. (D), relative fungal biomass, shown as the relative abundance of *Fulvia fulva* DNA (based on fungal actin gene), compared to tomato DNA (based on tomato actin gene), at 3dpi, 6dpi, 10dpi and 14dpi. Error bars are standard error (SE). The asterisks indicate statistically significant differences between KO plants and WT plants determined by student’s t-test (*p-value <0.05** p-value <0.01*** p-value <0.001**** p-value <0.0001). ns indicates no significant difference.

### Fungal tolerance mechanisms to α-tomatine contribute to virulence on *SlGAME4*-KO

Inoculations were performed with *B. cinerea* isolates B05.10 and M3a on leaves of *SlGAME4*-KO mutant plants. M3a was previously reported to infect MM poorly, due to the absence in its genome of the *BcTom*1 and *Bcgt*28a genes, that confer tolerance to α-tomatine (You et al., 2024). Alike on a wild type MM host, isolate M3a caused a low incidence of expanding lesions on *SlGAME4*-KO plants which accumulate uttroside B instead of α-tomatine (Figure 6A). We then studied the role of fungal tolerance mechanisms, mediated by α-tomatine-responsive genes, in the virulence on *SlGAME4*-KO. A B05.10 mutant in which *BcTom*1 was deleted showed significantly reduced fungal virulence. Conversely, the overexpression of three distinct types of tomatinase genes including *BcTom1*, *SlTom1* and *CfTom1* as well as the glycosyltransferase gene *Bcgt*28a promoted M3a infection on *SlGAME4*-KO (Figure 6).

**Figure 6.**
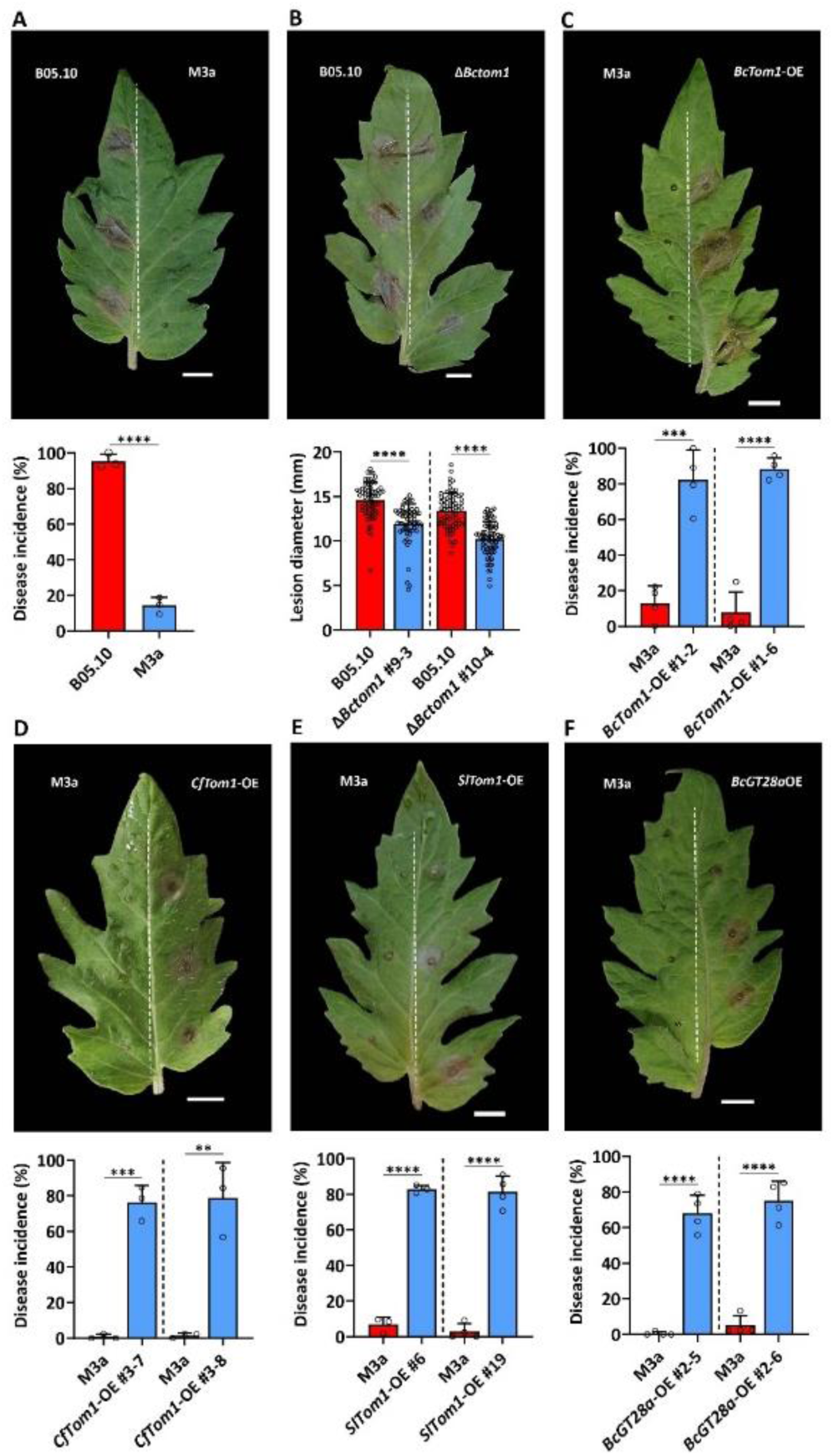
The role of fungal genes conferring tolerance mechanisms to α-tomatine in virulence on *SlGAME4*-KO. Disease incidence refers to the percentage of expanding lesions; lesion diameters were measured 3 days post inoculation on *SlGAME4*-KO leaves. A, Disease incidence of B05.10 compared with M3a. B, lesion diameters of wild type B05.10 compared with two *BcTom1*-KO mutants Disease incidence of wild type M3a compared with three tomatinase gene overexpression transformants *BcTom1*-OE (C) *CfTom1*-OE (D) *SlTom1* (E) and overexpression of glycosyltransferase gene *BcGT28a* (F). The asterisks indicate statistically significant differences determined by student’s t-test (** p-value <0.01*** p-value <0.001**** p-value <0.0001). Scale bar indicates 1 cm.

### BcTom1 contributes to virulence of *B. cinerea* isolate B05.10 on *Solanum nigrum*

Since uttroside B is the main saponin in *Solanum nigrum* (black nightshade), we characterized the importance of tolerance to α-tomatine in the interaction between *S. nigrum* and *B. cinerea*. As a negative control, we analysed the interaction with potato (*S. tuberosum*) which also produces alkaloid saponins, however with a different oligosaccharide chain. We first analyzed the expression profile of α-tomatine-responsive genes after inoculation on leaves of *S. nigrum* and potato. Expression of the tested genes was strongly upregulated at 12 h after inoculation on *S. nigrum* (Figure 7A), whereas most genes exhibited unaltered transcript levels at any time point after inoculation on *S. tuberosum* (Supplementary Figure S3). M3a formed significantly smaller lesions than B05.10 (Figure 7B), while the B05.10 mutant in which *BcTom*1 was deleted showed significantly reduced virulence on *S. nigrum* (Figure 7C).

**Figure 7.**
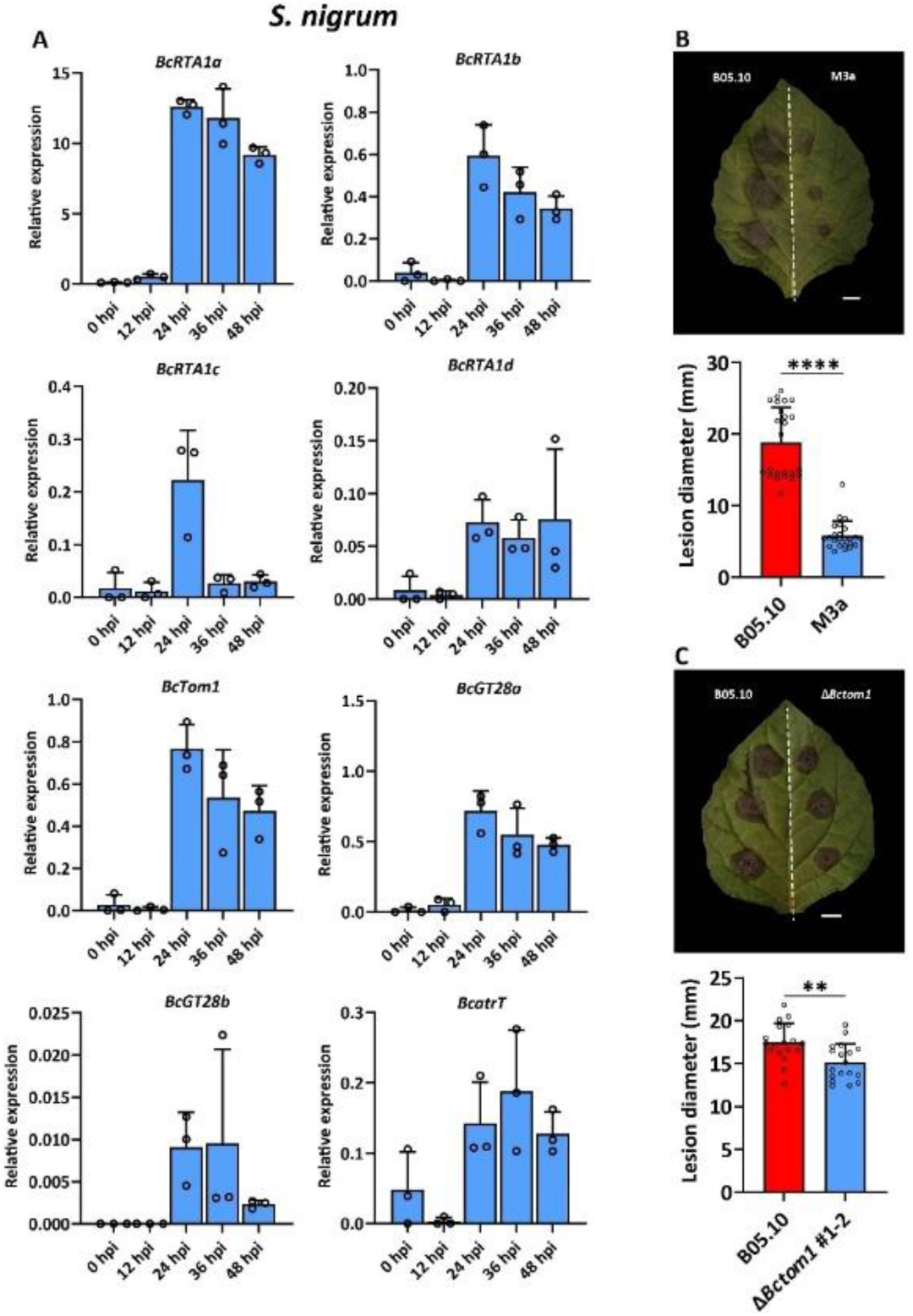
Characterization of interaction between *B. cinerea* and *S. nigrum*. Relative expression of α-tomatine-responsive genes in *B. cinerea* during infection on leaves of *S. nigrum* (A). Lesion diameters of B05.10 compared with M3a (B) and a *BcTom1*-KO mutant (C) as measured at 3 dpi. The asterisks indicate statistically significant differences determined by student’s t-test (** p-value <0.01**** p-value <0.0001). Scale bar indicates 1 cm.

## Discussion

In order to study the contribution of α-tomatine to basal resistance of tomato to microbial pathogens, we generated knockout lines in two different genes, *SlGAME4* and *SlGAME2*, which were reported by Itkin et al. (2013) to catalyze the first and the last dedicated steps of the α-tomatine biosynthetic pathway. Four independent homozygous *SlGAME4*-KO lines indeed lost the ability to produce α-tomatine, as well as tomatidine. Unexpectedly, all four independent homozygous *SlGAME2*-KO lines produced normal levels of α-tomatine. Deletions in the coding sequence of *SlGAME2* were close to the start codon and all caused a frameshift that should abolish the production of GAME2 protein in homozygous mutant lines. The observation that *SlGAME2-*KO plants accumulated normal levels of α-tomatine demonstrates that *Sl*GAME2 is not essential for α-tomatine biosynthesis in tomato. Itkin et al. (2013) described that *SlGAME2* clusters in the tomato genome on chromosome 7 with three other glycosyltransferase genes (*SlGAME1*, *SlGAME17*, *SlGAME18*) that are reported to be responsible for the first three steps in the glycosylation cascade of tomatidine. The tomato *Sl*GAME1/17/18/2 cluster is highly syntenic with the potato SGA biosynthetic cluster (Itkin et al., 2013). The protein sequence of SlGAME2 is orthologous to potato SGT3, which catalyzes the transfer of UDP-rhamnose (a hexose) to generate the potato SGAs α-solanine and α-chaconine (Itkin et al., 2013). On the contrary, SlGAME2 is supposed to use UDP-xylose (a pentose) as the donor-substrate. Evidence of the existence of β_1_-tomatine rhamnoside in tomato is lacking, despite numerous studies that have analyzed the SGA profiles in tomato. The fact that *SlGAME2-*KO plants still produce α-tomatine and that SlGAME2 is homologous to a potato rhamnosyltransferase thus questions the functional annotation of the *SlGAME*2 gene product proposed by Itkin et al. (2013). The tomato genome contains multiple glycosyl transferase genes, but none of these shows significant levels (>50%) of protein sequence identity to SlGAME2. Furthermore, transcripts of *SlGAME2* were only detected in young, developing vegetative tissues of tomato but were undetectable in adult expanded leaves, both by RT-qPCR and RNAseq (**Supplementary Figure S2** and **unpublished data**). By contrast, transcript levels of the other three genes that mediate the first steps of glycosylation of tomatidine (*SlGAME1*, *SlGAME17*, *SlGAME18*), are relatively high in all tissues. We therefore conclude that *SlGAME2* is not essential forα-tomatine biosynthesis. A glycosyltransferase was identified in *S. commersonii* that catalyzes attachment of xylose to triose SGAs, and thereby confers resistance to Colorado potato beetle and *Alternaria solani* (Wolters et al., 2023). A recent study by Gharat et al. (2025) showed that the tomato gene Solyc12g009930, the ortholog of the *S. commersonii* xylosyltransferase gene, indeed mediates the transfer of xylose to β1-tomatine and thereby completes the synthesis of α-tomatine. We hypothesize that knocking out the Solyc12g009930 gene in tomato will not only abolish the production of α-tomatine, but would also affect the last step of uttroside B synthesis, as it shares an identical tetrasaccharide moiety with a terminal xylose. It can be anticipated that a failure to add the terminal xylose would result in a strong reduction of fungitoxicity of uttroside B, analogous to the loss of toxicity that is exhibited by β1-tomatine.

The observed up-regulation of α-tomatine-responsive *B. cinerea* genes during infection on *SlGAME4*-KO leaves, which do not contain α-tomatine, suggested the presence of other inducers. The LC-MS analysis confirmed that *SlGAME4*-KO plants accumulate steroidal saponin uttroside B instead of steroidal glycoalkaloids (Figure 3 and S1). We previously reported that the expression of α-tomatine-responsive genes can also be induced by the steroidal saponin digitonin from *Digitalis purpurea*, which contains a pentasaccharide moiety that structurally resembles the tetrasaccharide moiety of α-tomatine and also possesses fungitoxic activity by permeating fungal membranes (You et al., 2024). Interestingly, uttroside B also contains four sugar moieties including terminal xyloside and glucoside in the lycotetraose unit (Grzech et al., 2024). The chemical similarity between α-tomatine makes it plausible to assume that uttroside Balso serves as target of fungal tomatinase, and can similarly induce expression of α-tomatine-responsive genes in *B. cinerea* during infection. This is supported by the observation that α-tomatine-responsive genes was observed during *B. cinerea* infection on *SlGAME4*-KO mutant and *S. nigrum* which both accumulate uttroside B (Figure 4 and 7A). Moreover, we showed that fungal tomatinases are still required for *B. cinerea* to successfully infect *SlGAME4*-KO and *S. nigrum* (Figure 6 and 7B).

Recent study has shown that uttroside B plays an important role in plant defense against insect pests (Boccia et al., 2024). In our study, change of tomato saponin compositions through KO of *SlGAME4* only slightly increased susceptibility to *B. cinerea* but did compromised resistance against other fungal pathogens including *<u>A. solani</u>*. *V. dahliae* and *F. fulva* (Figure 5). This observation indicates that uttroside B can largely replace the defensive role of α-tomatine in protecting plants from fungal pathogens. Besides, it is well-known that contents of steroidal glycoalkaloids such as α-tomatine strongly decreased during fruit ripening which is also seen in our study (Figure 3) (Bai et al., 2025; Kazachkova et al., 2021; Nakayasu et al., 2020; Sonawane et al., 2023). This process may compromised chemical defense in tomato mature fruits. Interestingly, *SlGAME4*-KO accumulate uttroside B instead of α-tomatine in green fruitsd and content of uttroside B did not strongly decrease in mature fruits (Figure 3). The presence of uttroside B in mature fruits might protect tomato from pathogens during postharvest. The secretion of α-tomatine from roots was reported to influence the tomato rhizosphere microbiome (Trivedi et al., 2020; Nakayasu et al., 2021). Therefore, it is interesting to characterize the effect of changing tomato saponin composition on manipulation of rhizosphere microbiome and potential influence of plant fitness using *SlGAME4*-KO plants in future studies.

## Experimental Procedures

### Generation of knockout lines in tomato

The *GAME* mutant lines were generated using a CRISPR/Cas9 genome editing strategy as described in Hanika et al. (2021). In brief, four sgRNAs were designed to target each gene (**Supplementary Table S1**). Transformation was carried out according to Huibers et al. (2013). T0 generations were screened and the lines carrying mutations in the coding sequences of target genes were used for seed production. Homozygous mutants were identified in T1 generation and their seeds were used for large scale experiments.

### Inoculation assays

Tomato plants used for fungal inoculations were grown at 20°C during daytime (16 h) and 19°C at night (8 h) with 60% humidity. Leaves from 5 to 6-weeks-old tomato plants were used for inoculation. Inoculation with *B. cinerea* was performed as described by You et al. (2023). Spores were suspended at 1x10^6^ spores/mL in 3g/L Gamborg’s B5 basal salt mixture supplemented with 15 mM sucrose and 10 mM potassium phosphate, adjusted to pH6.0. Six 2µL-droplets were inoculated on one leaflet and lesion diameters were measured at 3 dpi.

*A. solani* strain (altNL03003/CBS 143772) was grown on PDA plates and spores were collected as described by Wolters et al. (2019). Spores were suspended at 1x10^5^ spores/mL in Potato dextrose broth (PDB, 12g/L) supplemented with 0.3% agar. Six 10 µL-droplets were inoculated on one leaflet and lesion diameters were measured at 5 dpi.

*F. fulva* inoculation assays and fungal biomass quantification were performed as in Ökmen et al. (2013). Spores of *F. fulva* (race 0WU; CBS131901) were resuspended at 2x10^6^ spores/mL in tap water. Leaves of four- to five week old plants were sprayed on the lower side with spore suspensions. Eight plants per line were inoculated with *F. fulva*, while two plants per line served as uninoculated control group. Leaf samples were taken at 3 dpi, 6 dpi, 10 dpi and 14 dpi, freeze-dried and used for DNA isolation. qPCR was performed to assess the ratio of plant DNA to fungal DNA, using primers for amplifying the fungal *Ffactin* gene or the plant *Slactin* gene (primers in Supplementary Table S1).

*V. dahliae* inoculation assay was performed as described in Santhanam et al. (2013). Spores from *V. dahliae* strain JR2 were resuspended at 1x10^6^ spores/mL in demi water. Roots of 10-day-old tomato seedlings were dip inoculated in *V. dahliae* spore suspension. Eight to ten plants per line were inoculated with JR2, and two to five plants per line served as uninoculated control group. Canopy areas were measured at 15 dpi.

### Metabolite analysis and quantification of glycoalkaloids by LC-QqQ-MS

α-Tomatine and tomatidine concentrations were analysed in distinct tissues including roots, stems, young leaves, mature leaves, green or red ripe fruit in 4-5 biological replicates. Frozen tissue samples were freeze-dried, ground into powder and 5 mg from each sample was used for extraction in methanol containing 0.1% formic acid using 15 min sonication. After centrifugation, supernatant was filtered and used for LC-QqQ-MS analysis as described in You et al. (2024).

### Metabolite analysis by UHPLC–HRMS

Metabolites were analyzed by ultra-high-performance liquid chromatography-high-resolution mass spectrometry (UHPLC–HRMS). Analyses were performed using a Vanquish™ Horizon UHPLC system interfaced with an Exploris™ 120 Orbitrap mass spectrometer (Thermo Fisher Scientific, Dreieich, Germany). Frozen leaf and fruit tissue samples were freeze-dried, ground into powder and 5 mg from each sample was used for extraction in methanol containing 0.1% formic acid using 15 min sonication. For each sample, 5 µL of extract was injected onto an Acquity UPLC BEH C18 column (1.7 µm particle size, 2.1 × 150 mm; Waters) maintained at 40 °C. Chromatographic separation was conducted at a constant flow rate of 400 µL min⁻¹ using a binary mobile phase system consisting of 0.1% (v/v) formic acid in water (mobile phase A) and 0.1% (v/v) formic acid in acetonitrile (mobile phase B). The gradient elution program was as follows: isocratic 5% B from 0.0 to 1.0 min; linear increase to 75% B from 1.0 to 22.0 min; linear ramp to 90% B from 22.0 to 23.0 min; isocratic hold at 90% B from 23.0 to 26.0 min; linear ramp to 5% B from 26.0 to 27.0 min followed by re-equilibration at 5% B from 27.0 to 30.0 min.

For compound putative annotation, samples were analyzed using combined full-scan MS and data-dependent MS/MS acquisition (Full MS/dd-MS², top-n), separately in negative- and positive-ionization modes. Full-scan MS data were acquired over an m/z range of 90–1350 at a resolving power of 60,000 (m/Δm at m/z 200), an automated gain control (AGC) target of 1 × 10⁶, and a maximum injection time of 100 ms. The ESI source parameters were set to a spray voltage of 3.0 kV (negative-ionization mode) and 2.5 kV (positive-ionization mode), and a capillary temperature of 290 °C. Data-dependent MS² spectra were acquired for the four most intense precursor ions per scan at a resolving power of 15,000 (m/Δm), using an AGC target of 1 × 10⁵, a maximum injection time of 118 ms, and an isolation window of 1.0 m/z. Fragmentation was achieved using stepped normalized collision energies (NCE) of 20, 40, and 100 eV (%) to generate diagnostic fragment ions supporting the annotation.

### Gene expression analysis

Total RNA was extracted from different tissues of tomato, potato or *S. nigrum* using Maxwell™ 16 RNA Purification Kits (Promega). Relative expression of fungal genes was quantified by RT-qPCR according to Qin et al. (2023), using primers described in **Supplementary Table S1**. The transcript levels of two *B. cinerea* housekeeping genes *BcTUBA* (Bcin01g08040) and *BcSMT3* (Bcin11g03430) were used for gene expression normalization

## Acknowledgements

The authors acknowledge Bert Essenstam (Unifarm, Wageningen UR) for excellent transgenic plant care and Ioannis Thanos (MSc) for performing a part of the qRT-PCR analysis. We acknowledge the support by Dr. Carlos Sanchez Arcos and Bert Schipper (Wageningen UR) in performing LC-MS2 runs of plant samples, and by dr. Marianna Boccia and dr. Sarah O’Connor (Max Planck Institute for Chemical Ecology, Jena, Germany) who provided an aliquot of uttroside B as LC-MS standard.

---

**Supplementary Figure S1.**
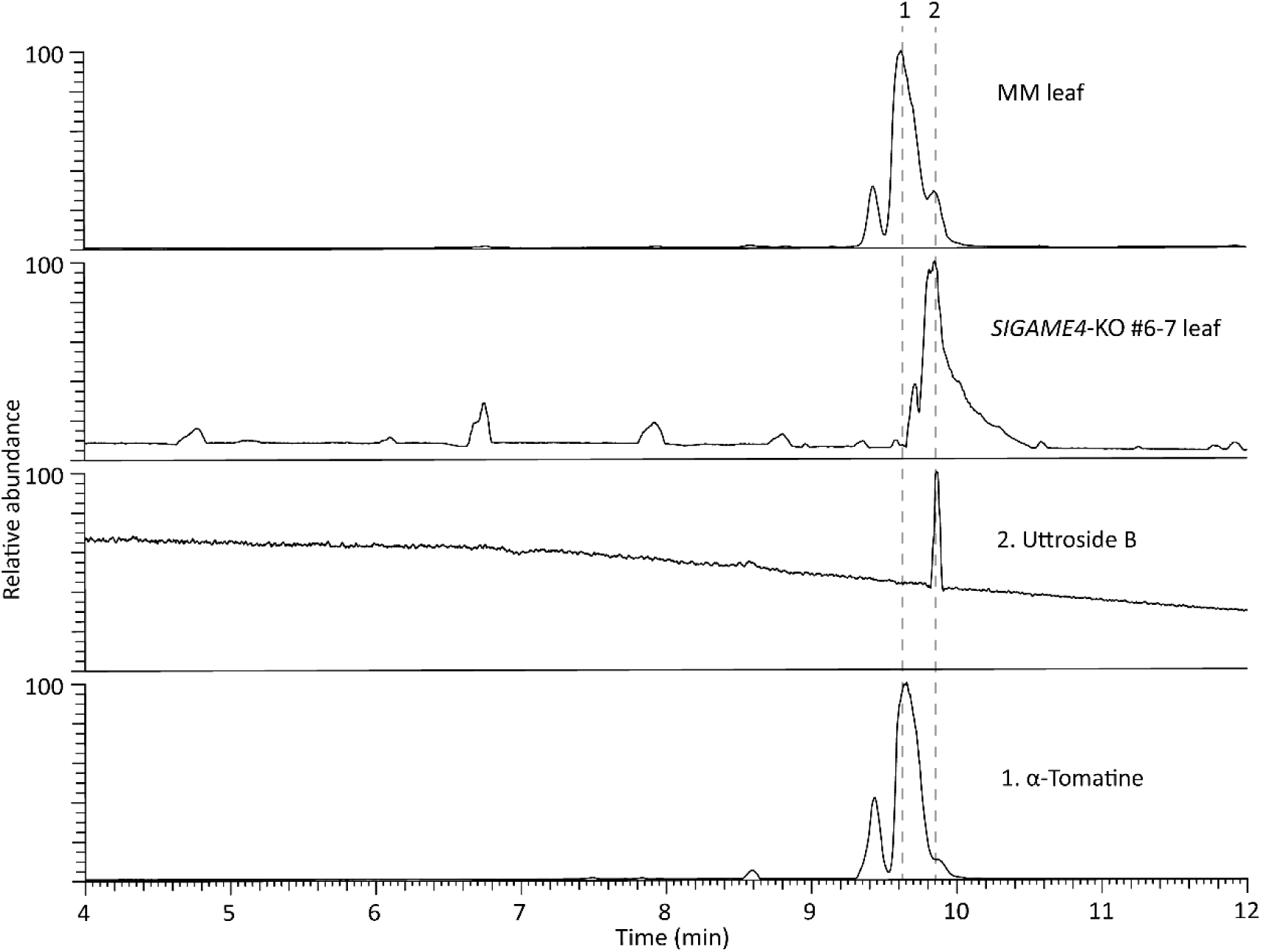
LC-MS/MS analysis of leaves from wild type Moneymaker (A) and *SlGAME4*-KO mutant genotypes (B). Standards for uttroside B and α-tomatine are shown in the panels C and D. The profile of MM is shown for one leaf and was representative for that of leaves from three other plants. The profile of the mutant shown here is for *SlGAME4*-KO #6-7, and was representative for that of leaves from the independently generated mutant genotypes *SlGAME4*-KO #8-1, *SlGAME4*-KO #12-2 and *SlGAME4*-KO #25-2.

**Supplementary Figure S2.**
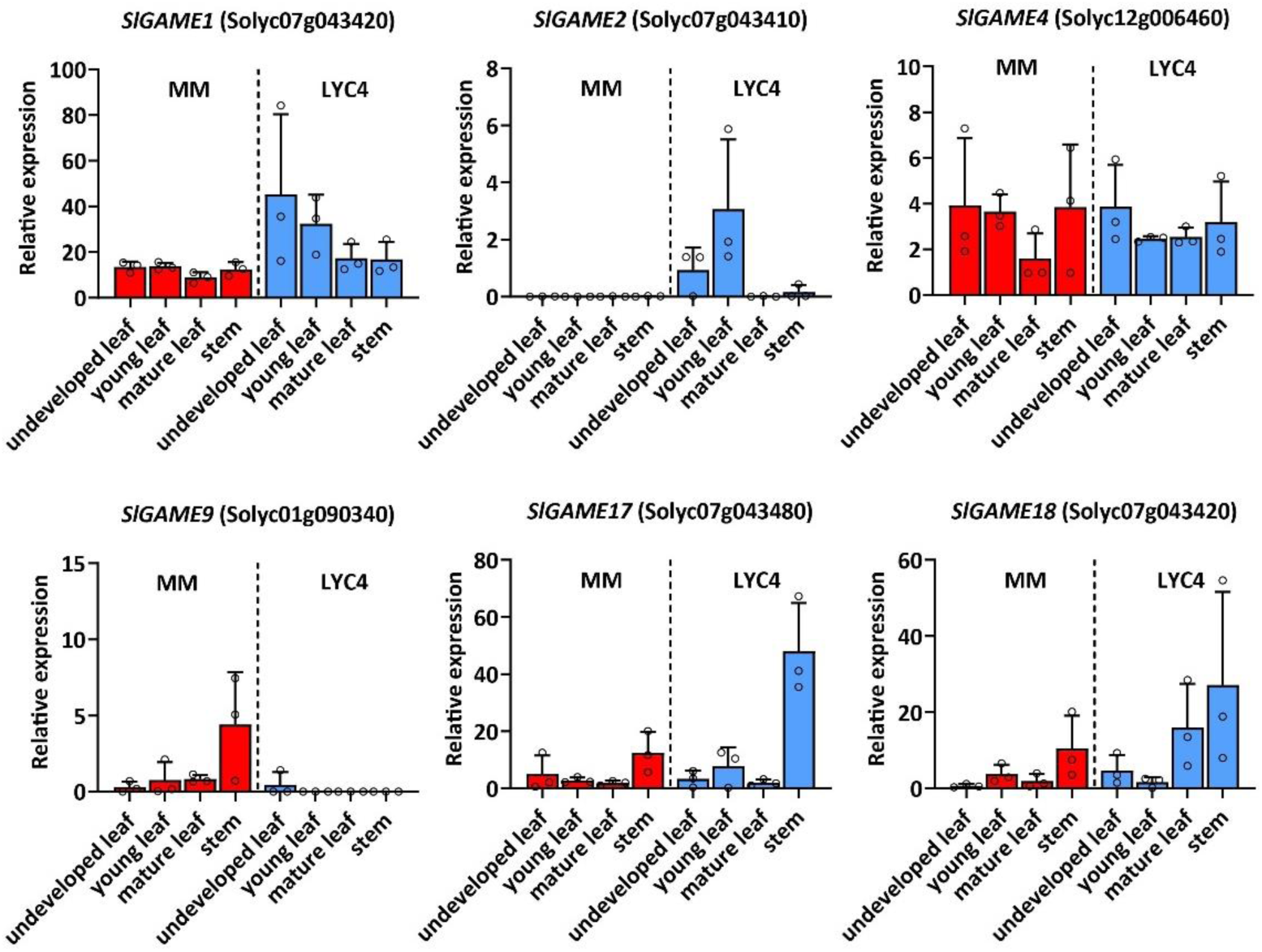
Relative expression of *GAME* genes in different tissues from *S. lycopersicum* MM and *S. habrochaites* LYC4 analyzed by RT-qPCR. Two house-keeping genes *CAC* and *GAPDH* were used as references. Error bars are standard error (SE) of three biological replates.

**Supplementary figure S3.**
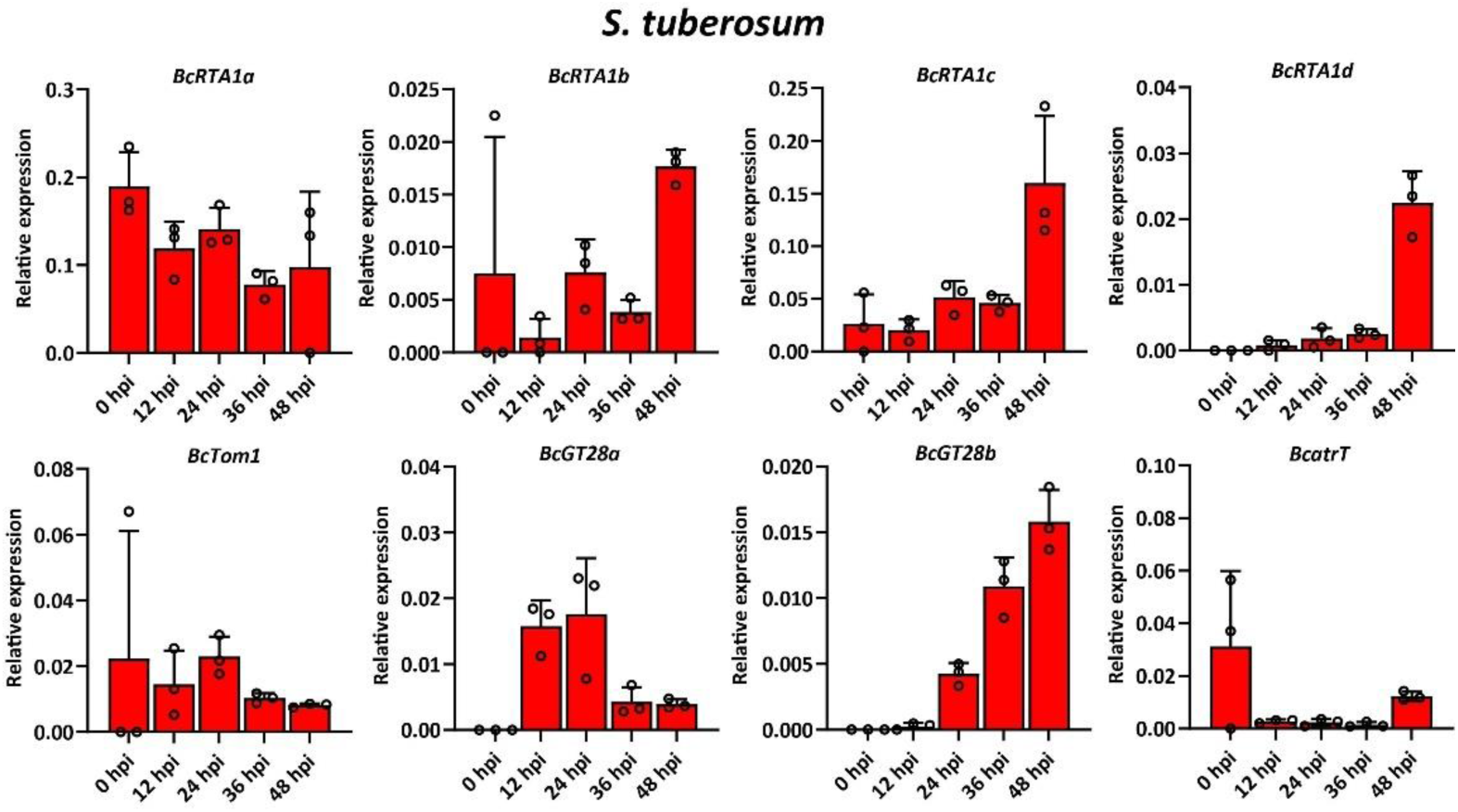
Relative expression of α-tomatine-responsive genes in *B. cinerea* during infection on leaves of *S. tuberosum.* Error bars are standard error (SE) of three biological replates.

**Supplementary Table S1.**
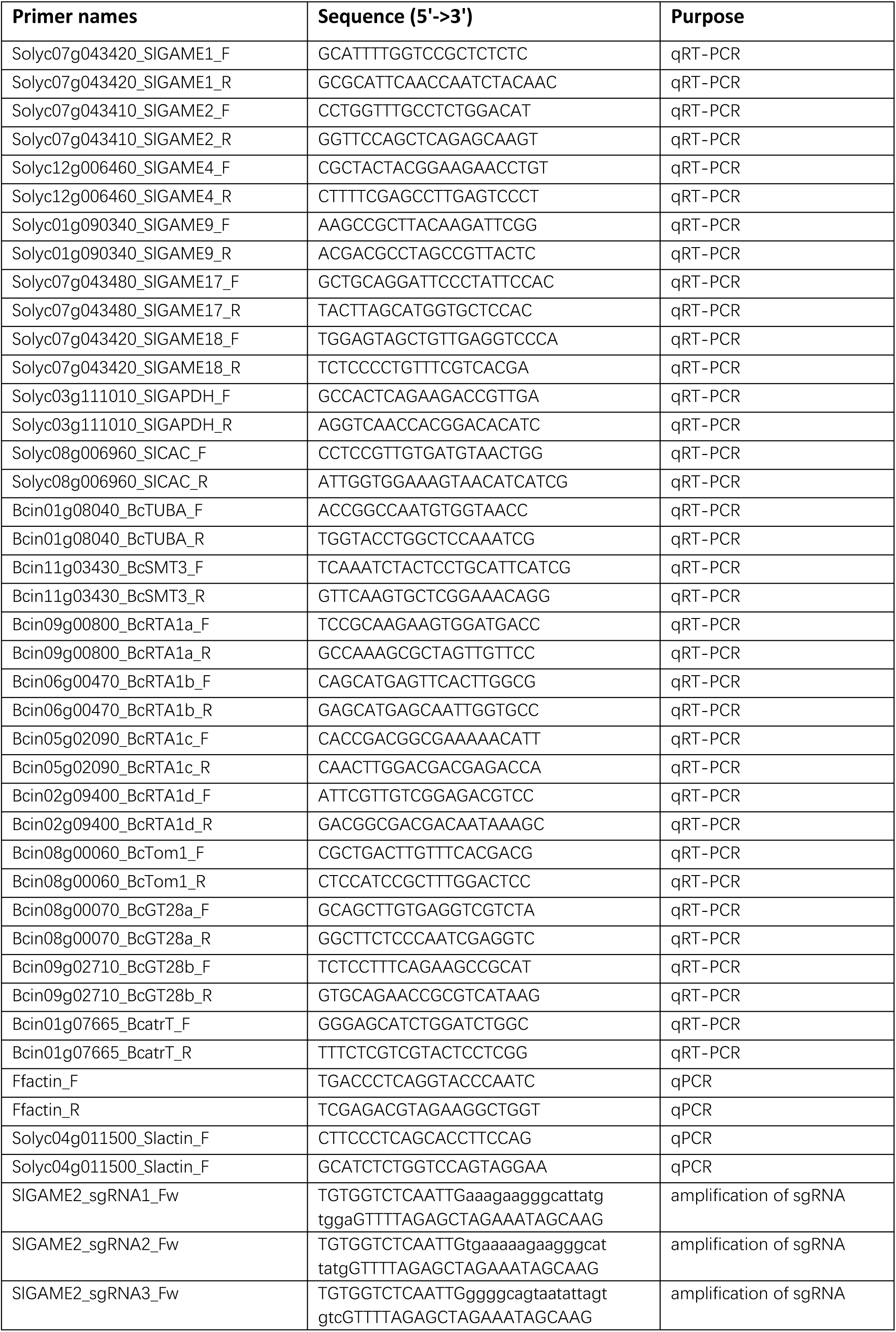
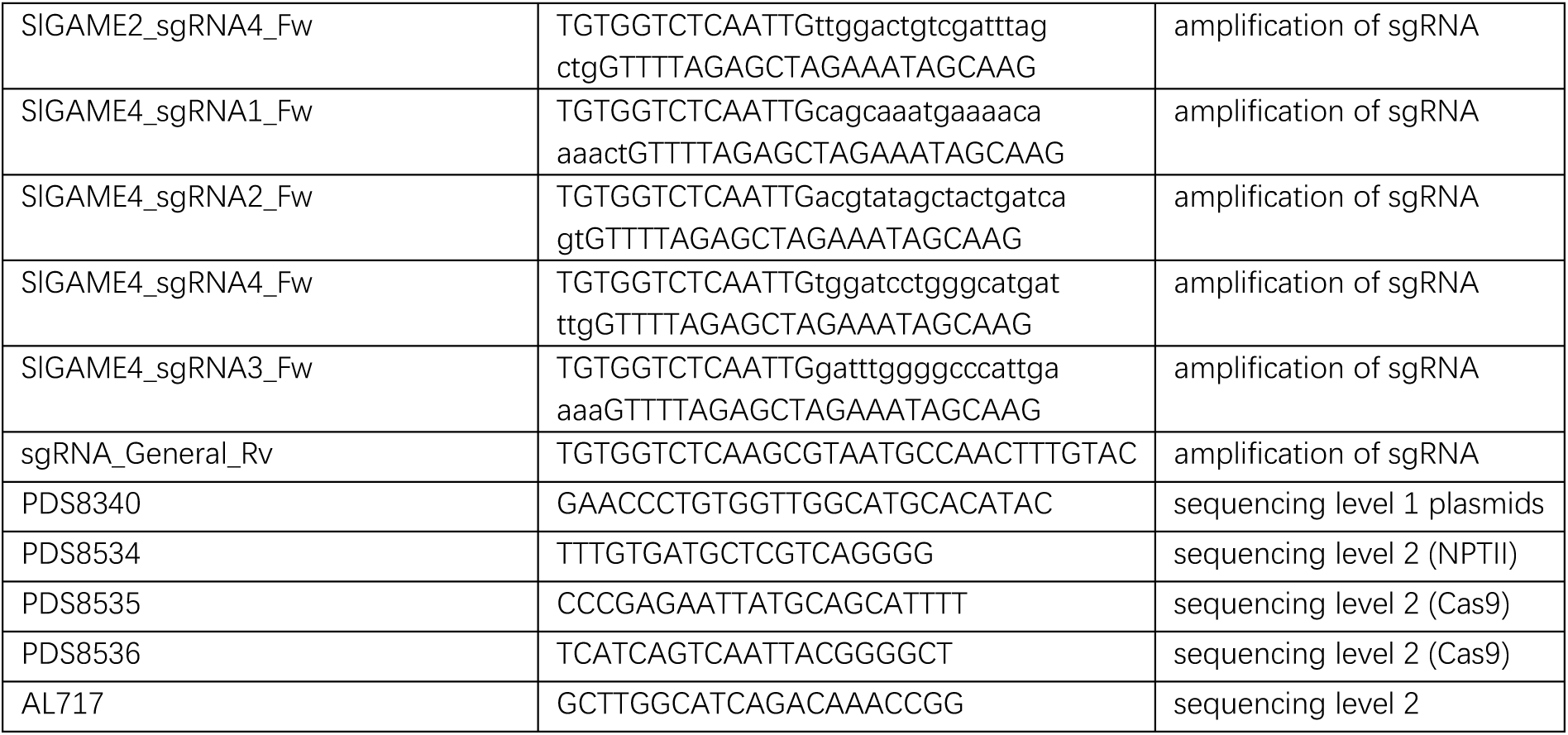
Primers used in this study.

## Notes

### Competing Interest Statement

The authors have declared no competing interest.

